# Phase of firing does not reflect temporal order in sequence memory of humans and recurrent neural networks

**DOI:** 10.1101/2022.09.25.509370

**Authors:** Stefanie Liebe, Johannes Niediek, Matthijs Pals, Thomas P. Reber, Jenny Faber, Jan Bostroem, Christian E. Elger, Jakob H. Macke, Florian Mormann

## Abstract

A prominent theory proposes that the temporal order of a sequence of items held in memory is reflected in ordered firing of neurons at different phases of theta oscillations ^1^. We probe this theory by directly measuring single neuron activity (1420 neurons) and local field potentials (LFP, 921 channels) in the medial temporal lobe of 16 epilepsy patients performing a working memory task for temporal order. We observe theta oscillations and preferential firing of single neurons at theta phase during memory maintenance. We find that - depending on memory performance - phase of firing is related to item position within a sequence. However, in contrast to the theory, phase order did not match item order. To investigate underlying mechanisms, we subsequently trained recurrent neural networks (RNNs) to perform an analogous task. Similar to recorded neural activity, we show that RNNs generate theta oscillations during memory maintenance. Importantly, model neurons exhibit theta phase-dependent firing related to item position, where phase of firing again did not match item order. Instead, we observed a mechanistic link between phase order, stimulus timing and oscillation frequency - a relationship we subsequently confirmed in our neural recordings. Taken together, in both biological and artificial neural networks we provide validating evidence for the role of phase-of-firing in memory processing while at the same time challenging a long-held theory about the functional role of spiking and oscillations in sequence memory.

## Introduction

How do we remember the temporal order of a sequence of events? Performing this kind of task is an integral part of our ability to encode, maintain and retrieve memories within their spatial and temporal context^2^. The medial temporal lobe (MTL) has been heavily implicated in memory processing at the neural level. A prominent finding is that neurons within MTL regions, such as the hippocampus, exhibit elevated, stimulus-specific spiking activity during the maintenance period of working memory tasks^3–7^. Another hallmark signature of neural activity within the MTL are oscillations in the frequency range of 2-8 Hz, commonly known as the theta-band. Theta oscillations can be measured from LFP or intracortical EEG/ECoG and have been ubiquitously observed in many species during memory processing^8^. Specifically, the amount of oscillatory activity, i.e. theta power, increases during memory maintenance and correlates with memory load and task performance^9–11^.

Combined measurements of spiking and LFPs from MTL have further established an important link between the single neuron firing and theta oscillations: Firing of MTL neurons depends on theta phase - so called spike-phase coupling - and phase of firing contains information about multiple spatial locations during sequential spatial encoding as well as spatial memory tasks in rodents^8;12–15^.

In analogy to spatial memory in rodents, it has been suggested that this so-called ‘temporal code’ is also suitable to represent multi-item sequences during working memory of non-spatial information^16^. Specifically, a prominent computational model by Lisman and collegues hypothesizes that the order of items held within memory is represented by spiking of sequentially re-activated neurons at different phases of theta-oscillations^1;17^.

Indeed, human MTL neurons show preferential firing with respect to theta phase in memory tasks^18^, the magnitude of spike-phase coupling is predictive of subsequent memory performance^19^ and spiking relative to theta phase contains non-spatial information, namely stimulus identity^20;21^. However, thus far it remains unclear how memorizing the sequential order of multiple items is implemented at the neural level in human MTL. In particular, it is unknown, whether a sequence of memorized items is associated with, first, differences in theta-related phase of firing of single neurons and, second, whether the order of phase-of-firing matches the item order - as hypothesized by Lisman’s theory^1^.

Here, we sought to answer these questions by directly measuring both spiking of MTL neurons and LFPs while participants had to maintain the temporal order of a sequence of items within working memory. We also investigated potential underlying neural mechanisms by training recurrent neural networks (RNNs) to perform an analogous task, without explicitly instructing RNNs on *how* to solve it. Our results show emerging theta oscillations and spike phase-coupling in both recorded and modeled neural activity during working memory, where phase of firing is related to item position within a sequence. Surprisingly, however, phase order did not match item order, in contrast to Lisman’s theory. Instead, our modeling suggests that phase-order could arise as a function of inter-stimulus interval and oscillation frequency - a relationship we subsequently confirmed in our neural recordings. Our findings thus validate, but also challenge a long-standing theory about the role of spiking and oscillations in memory function.

## Results

### No effect of item position on spike rate during memory

We recorded spiking activity of 1420 units and LFPs from 921 channels in medial temporal lobe regions including the hippocampus (HPC), enthorinal cortex (EC), parahippocampal cortex (PHC) and amygdala (A) in 16 chronic epilepsy patients undergoing invasive seizure monitoring for presurgical evaluation. Patients performed a sequential multi-item working-memory task as illustrated in **Fig. 1 a**. After a fixation period, four randomly chosen pictures out of a set of 8 were sequentially presented to the patients for 200 ms each with a 400 ms time difference between each stimulus onset. Stimulus presentation was followed by a delay period of 2500 ms (+/- 100 ms) after which a panel comprising four rows of possible picture sequences appeared. One of them matched the previously presented sequence, and patients indicated a match by pressing the associated row number on a computer keyboard. Each patient performed markedly above chance but still in a range that allowed us to compare neural activity for correct vs. incorrect trials later on. Moreover, median reaction time (RT) showed a typical negative correlation with mean performance across subjects (**Fig. 1 b**). Thus, our behavioral results indicate that subjects understood the task well and were generally attentive during participation.

**Figure 1.**
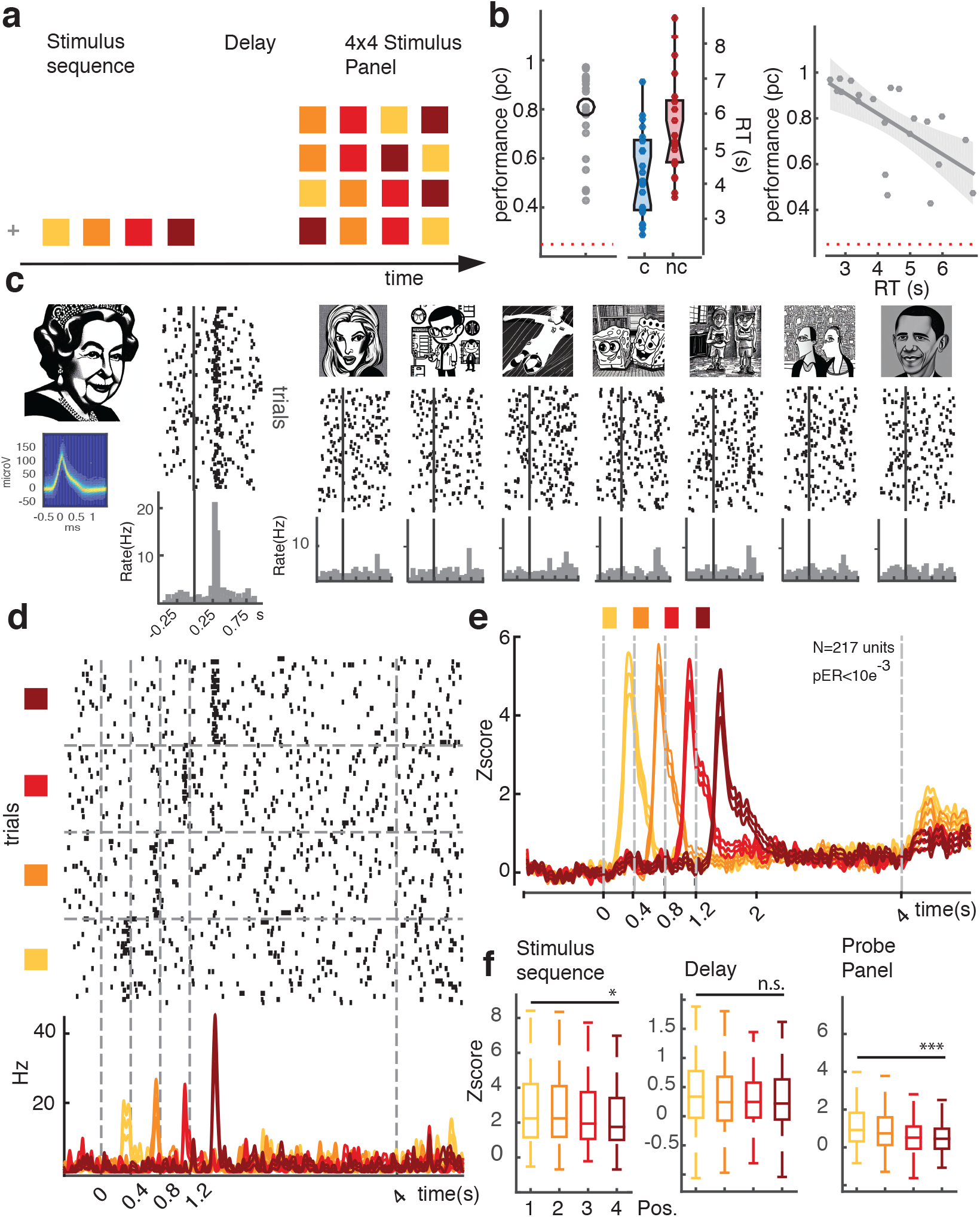
Experimental paradigm, behavioral performance and spike rate modulation. **a**. Experimental design of multi-item temporal-order working memory task. **b**. Left panel: Individual mean percentage correct per session (mean percentage correct 77%, SD 17,8%, for each patient compared to chance performance at 25% (red dashed line) Binomial test p<10^−3^), black circle: average across subjects) and median reaction times (RT) for correct (c) vs incorrect (nc) trials. Right panel: Median reaction times (RT) plotted against mean percentage correct with least-squares linear fit and s.e.m. shows negative correlation (Spearman rank-correlation coefficient -0.69, p<10^−3^, median RT 4514 ms, SD 1328 ms). **c**. Spike waveform density plot, raster plots and PSTHs from a hippocampal unit to all stimuli shown in one session. Vertical lines correspond to stimulus onset. This unit showed a significant rate increase to only one of the eight stimuli (PS, Wilcoxon signed-rank test p<10^−8^). **d**. Spiking activity of the same unit across the entire trial period in response to the PS shown at four different positions within the sequence (1-Way repeated measures ANOVA, F=3.05, p<0.05 during the stimulus period) **e**. Convolved PSTH and s.e.m. (Z-score relative to baseline) in response to the PS of each individual neuron, averaged across all stimulus-responsive units for the entire trial period (sequence position color-coded) **f**. Median spike rates for each stimulus position and specific trial periods. (Kruskal-Wallis non-parametric ANOVA on stimulus-evoked response, Chi2=8.34, p<0.01,Probe Panel Chi2=39.5, p<10^−4^, Delay Chi2=2.55, p>0.05, N=217). Note: (Stimuli used in the experiments can not be displayed according to the inclusion rules of biorarxiv, and have been replaced by thumbnails generated with stable diffusion, https://huggingface.co/spaces/stabilityai/stable-diffusion).

While spike rates of MTL units are typically modulated by stimulus identity, little is known about whether the sequence position of stimuli also affects their spiking in a systematic fashion. To address this question, we first identified 217 highly responsive units by comparing stimulus-evoked activity during encoding to a pre-stimulus baseline period (Wilcoxon signed-rank test across all positions, criterion of *α <* 10^−3^ for comparing stimulus vs. baseline interval, HPC:N=82, EC:N=25, PHC:N=55, A:N=55). We show a representative example for a highly stimulus-responsive unit that exhibited a significant rate increase for one of the eight stimuli, the so-called preferred stimulus (PS) in **Fig. 1 c**. When analyzing spiking in response to the PS at each of the four positions within the sequence (**Fig. 1 d**) stimulus-evoked activity of this unit differed significantly between item positions during encoding, with the largest evoked response at the end position of the sequence and no modulation of spike rate during delay or the probe period. When assessing spiking activity across all stimulus responsive units for the PS at each item position, however, we found that stimulus-evoked responses were typically largest for the first position and appeared to systematically decrease with increasing sequence position (**Fig. 1 e**). Interestingly, we observed a similar effect during the panel presentation. Here, spiking activity was largest whenever the units’ preferred stimulus had been shown as the first within the initially presented sequence, even though the visual input consisted of a 4×4 picture grid (**Fig. 1 f**). In contrast, during the delay period spike rates did not significantly differ between stimulus positions even though spiking was significantly elevated during the delay compared to the pre-stimulus baseline, suggesting active maintenance-related activity similar to previous findings^3^, (Wilcoxon signed-rank-test, Z>2.7, p<0.001 for each region).

In summary, MTL neurons show robust stimulus-specific encoding responses during visual presentation, where spiking decreases as stimulus position increases. These findings might be related to the so-called primacy effect that has been observed in serial memory paradigms for mesoscopic brain signals where stimuli shown at the first position within a sequence also elicit larger neural responses^22^, arguably due to increased allocation of attentional resources during encoding^23^. In contrast, we found no such differences during memory maintenance. Thus, even though participants were explicitly asked to remember the order of the encoded stimuli, this does not seem to be reflected in systematic spike-rate changes during memory in the MTL.

### Theta oscillations and spike-phase coupling during memory

Previous studies have observed increases in theta oscillations, i.e., power (2-8 Hz) during working memory maintenance^9;24^. Thus, we first asked whether we find similar enhancements in our task. An example time- frequency spectrogram recorded from the hippocampal site shown in **Fig. 1** is plotted in **Fig. 2 a**. A clear increase in theta power around 2.8 Hz can be seen, starting after stimulus presentation and extending throughout the delay (median power baseline vs. delay, Wilcoxon signed-rank test p<0.01). We generally observed a similar effect comparing power across all LFP channels of stimulus responsive units between baseline and delay (Wilcoxon signed-rank test N=217, Z=3.7, p<0.001, see also Extended Data **Fig. 1 a/b** S1). Comparing single areas, the overall increase was mainly present at hippocampal and entorhinal sites even though each region contained a significant proportion of channels exhibiting elevated theta during maintenance (HPC:40%, EC:48%, PHC:20%, A:36%, Binomial test p<10^−4^ at *α*=0.01). Thus, our results confirm earlier findings on memory-related power increases in theta during working memory and provide the basis for our following analyses.

**Figure 2.**
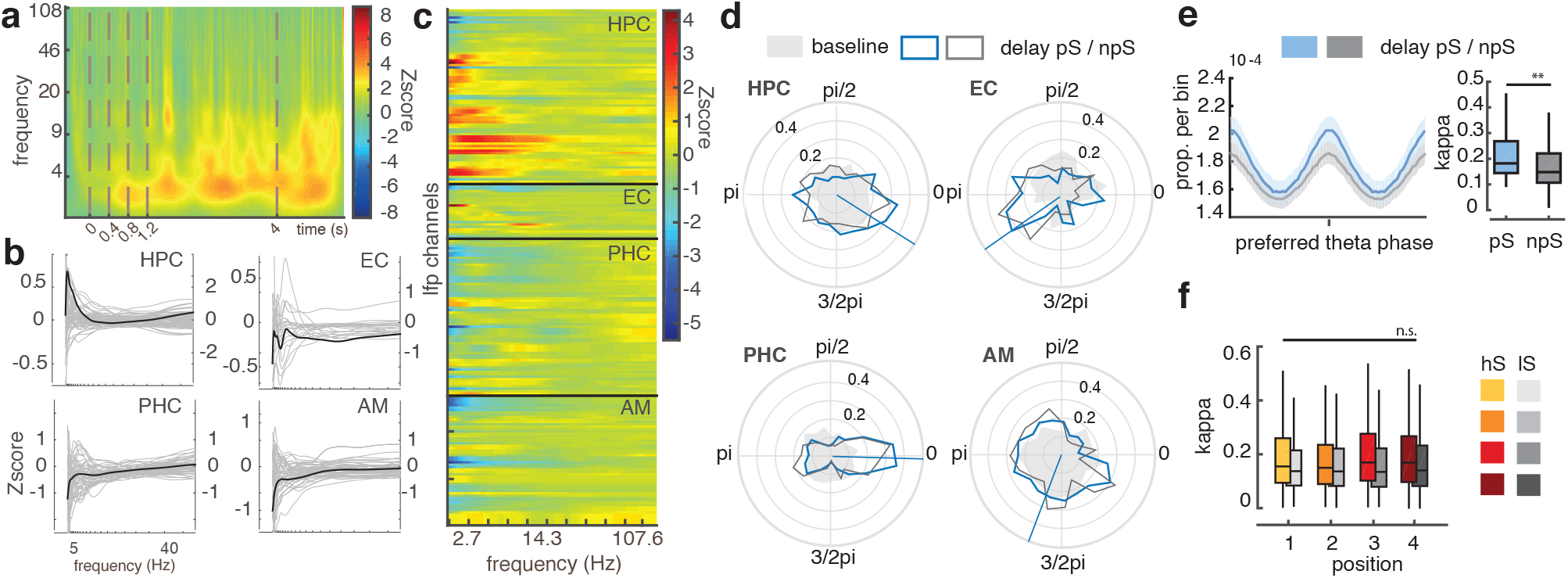
Theta oscillation and spike-phase modulation during sequence maintenance. **a**.Example of normalized time-frequency spectrum (power Z-score relative to baseline) in anterior hippocampus (same recording site as in 1) showing sustained increase in theta power during the delay. **b** Normalized power spectra during the delay per MTL region (black median, gray single channels). **c** Same as **b**, but color-coded for each channel. **d** Preferred phase distribution in theta range for all stimulus-responsive units during baseline (shaded) and delay where sequences contained the PS (blue) or not (gray), lines show mean preferred phase angle during delay. **e** Left panel: Mean (95% CIs) spike-phase histogram centered at each individual unit’s preferred phase during the delay, separately for PS vs. NPS trials. Right panel: Median estimated concentration parameters *κ* for PS vs. NPS trials based on individual van Mises fits. **f** *κ* separately per stimulus position for more selective (yellow-red) vs. less selective units (light-dark gray).

Combining single unit and LFP recordings, we next analyzed spiking as a function of theta phase. In each MTL region, we observed a non-uniform distribution of theta phases at firing, for which preferred phase angles differed considerably across MTL regions (non-parametric multi-sample test comparing median angles between regions P=50.9, p<10^−^10). Spike-phase coupling was already present during pre-stimulus baseline but significantly increased during memory maintenance at the population and individual units’ level (population-based Z-score Rayleigh test for N=217 units Z=9.7/3.2, p<0.01/p<0.05 for Delay/Baseline, respectively; for single areas: Z(HPC) = 4.32/4.1, Z(PHC)= 14.3/4.51, Z(AM)=10.58/3.89, Z(EC)=15.1/4.52, Delay all areas p<0.02/Baseline all p<0.04, Median comparison Rayleigh-based Z-Scores on individual units, Wilcoxon signed-rank test Z>9.63, p<10^−10^ for all regions, see also **Fig. S1 c**). Importantly, the effects were also observed when correcting for spike rate differences between baseline and delay (see Methods Section). To assess whether spike-coupling is associated with memorizing specific stimulus information, we computed mean spike-phase histograms from delay activity centered at each individual unit’s preferred phase, and computed the concentration parameter kappa (*κ*), derived from van Mises fits to spike-phase histograms, to assess strength of spike phase-coupling (**Fig. 2 e**, left/right panel, respectively). We observed that modulation of spiking at theta phase was significantly elevated during maintenance if the PS was encoded during the sequence, even after controlling for spike rate differences, which indicates that memory-related enhancement of spike-phase coupling is stimulus-specific (Wilcoxon signed-rank test comparing median *κ* values based on individual unit’s van Mises fits, N=217, Z=8.65, p<0.001). We next asked whether the magnitude of spike-phase coupling also depends on serial position of stimuli. To this end, we did not observe a difference between these conditions (**Fig. 2 f**, Kruskal-Wallis test based on individual *κ* values, p>0.05, Chi2<3.9). Taken together, maintenance of multiple stimuli correlated with increased spike phase-coupling across the population as well as for individual units. Specifically, phase-of-firing distributions became less uniform as units showed more similar phase preferences during the delay. For individual units, increased spike-phase coupling was specific to the encoded stimulus. Interestingly, similar to our spike rate analyses, the magnitude of phase coupling was not systematically related to the position of the stimulus within a sequence.

### Phase-of-firing depends on sequence position

In our next analyses, we focused on the temporal relationship between theta oscillations and spiking. We first asked whether the preferred phase of firing during the delay varies with item position in a sequence, as this is one of the central model predictions by^1^. When analyzing spike-phase histograms, we found that single units showed differences in preferred phase of firing depending on the sequence position of the PS (**Fig. 3**). To quantify this relationship, we computed the circular variance explained between phases at different sequence positions (*V*_*ex*_, see Methods). To assess statistical significance for each unit, we compared *V*_*ex*_ to a null distribution derived from randomly shuffling position labels and performing a permutation test against shuffled *V*_*ex*_ per unit (p<0.001, Permutation test for units shown in **Fig. 3**.

**Figure 3.**
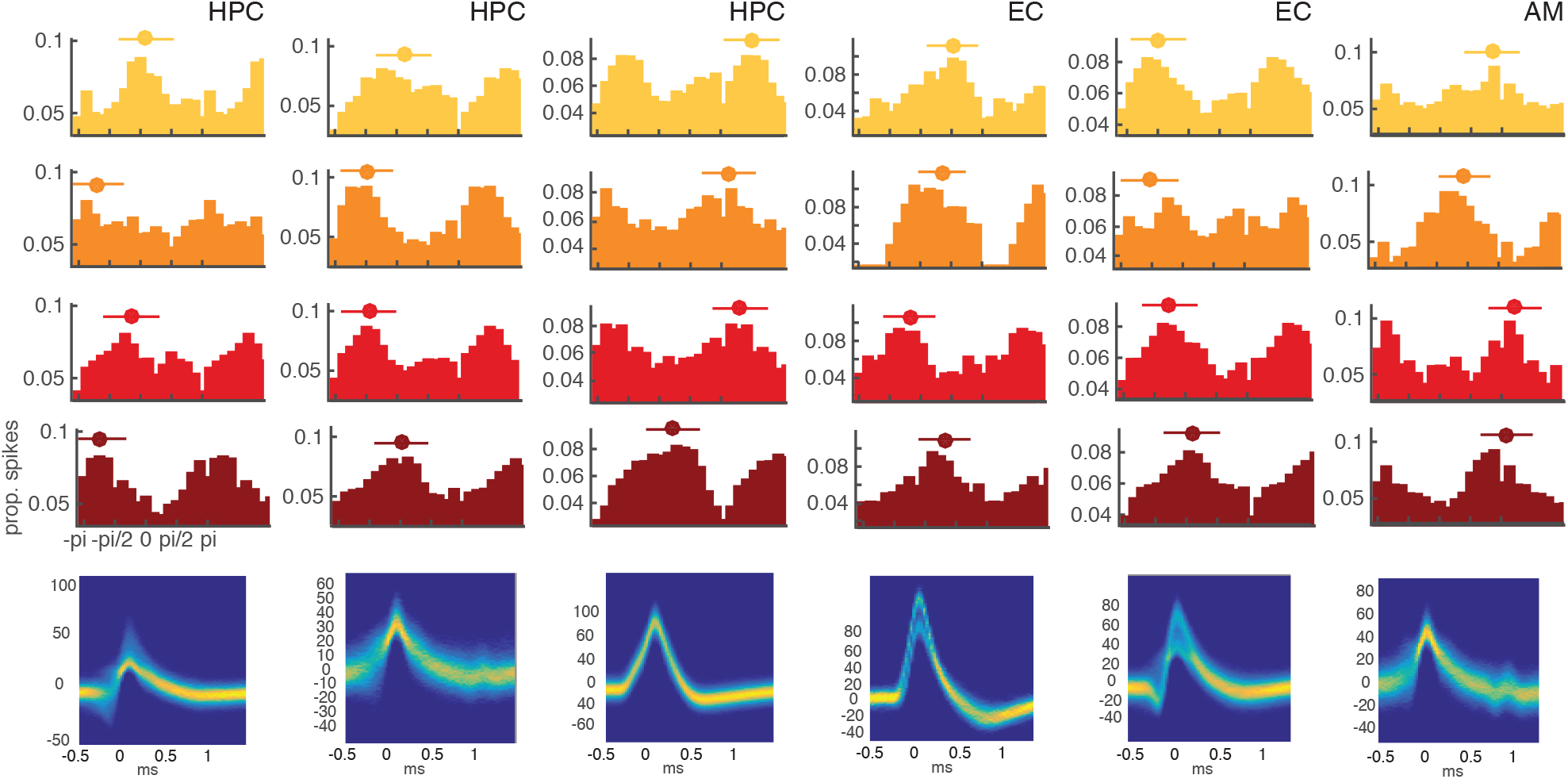
Phase of firing in single neurons encodes sequence position. Spike waveforms (bottom row) and spike-phase histograms (upper rows) of single units at different item positions (color coded for sequence position, HPC-hippocampus, EC-entorhinal cortex, AM-amygdala). Mean preferred phase and circular standard deviation are shown above the histograms in each plot.

Analyzing *V*_*ex*_ as a function of frequency for highly stimulus selective units, we observed phase differences between sequence positions across all investigated theta bands and different MTL regions **Fig. 4 a**. However, when we compared *V*_*ex*_ for shuffled vs. non-shuffled position labels, we found the largest differences in the lower theta frequency range (2-3 Hz), in line with earlier findings investigating spike-phase coupling with respect to stimulus identity^20^ (**Fig. 4 b**). To test whether this effect was related to task performance, we repeated the analysis separately for groups of correct and incorrect trials and found indeed that *V*_*ex*_ was significantly enhanced for correct, but not incorrect trials (**Fig. 4 c**, Permutation test non-shuffled vs. shuffled, p<0.05). Likewise, when directly comparing correct and incorrect trial-based estimates, *V*_*ex*_ was significantly larger, again for lower theta frequencies (Wilcoxon rank-sum test p<0.05 for 2/2.4 Hz). Critically, we observed a similar effect choosing only the one theta frequency per unit-LFP pair for which we observed the highest oscillatory power increase from baseline to delay (**Fig. 4 d**, paired T-test, T=2.23, p<0.02 across all units). This is important, as this analysis not only accounts for variance in oscillatory peaks between LFP channels^18^ and the number of statistical comparisons when analyzing multiple frequencies, but also uses an independent criterion for selecting a specific theta frequency.

**Figure 4.**
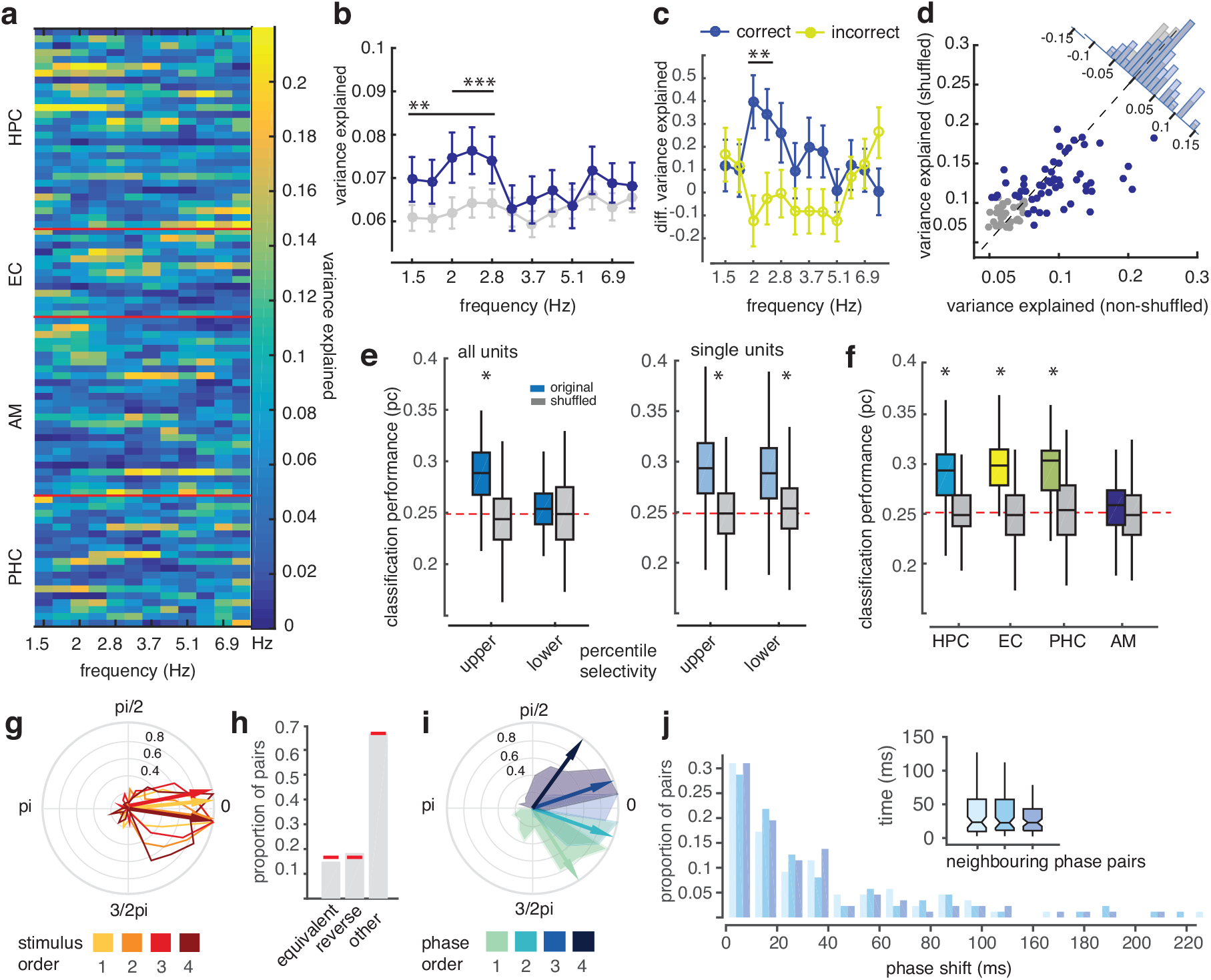
Preferred phase of firing varies with sequence position. **a**. Circular variance explained *V*_*ex*_ between positions as a function of frequency for units showing high stimulus selectivity (N=87, based on median split, units sorted by area). **b**. Average *V*_*ex*_ across units for non-shuffled (blue) and shuffled (gray) position labels as a function of frequency. Error bars denote s.e.m., stars indicate significance based on Permutation test p<0.01 (**) and p<0.001 (***), N permutations = 1999). **c**. Averaged difference in *V*_*ex*_ between non-shuffled and shuffled conditions for correct (blue) and incorrect (yellow) trials. **d**. Scatter plot of *V*_*ex*_ per unit shown for shuffled (y-axis) vs. non-shuffled position labels (x-axis) at the theta frequency with the respective highest power increase during the delay as well as corresponding group histogram (inset, blue dots correspond to values larger than upper 50th percentile across all units in either group). Decoding performance based on preferred mean phase of firing during delay across the population of units (left) and individual units (right) predicting one out of four different stimulus positions. Performance is plotted separately for high vs. low stimulus selective units on median split, and separately for non-shuffled and shuffled position labels. **f**. Decoding performance for different MTL regions. **g**. Phase of spiking histograms and mean direction across units shown in **a**. per stimulus position (color-coded). Plotted are circular differences in phase with respect to mean phase across all positions per unit **h**. Proportion of units for which phase of firing order for different positions is equivalent to item order, is reversed, or different. Red lines denote proportion of respective order expected by chance (1/6, 5/6 respectively). **i**. Equivalent plot as in **g**. except here phases are assigned a ‘position’ label based on recorded phase order, not stimulus order. **j**. Distribution of three consecutive pairwise phase shifts in ms based on neighboring phase groups shown in **g** as well as median pairwise time differences per neighboring pair.

In our next analyses, we tested whether sequence position can be decoded from phase of firing employing a support vector machine (SVM) algorithm. **Fig. 4 e** depicts classification performance (percent correct, PC) based on population activity (all units) and individual units, using shuffled position labels as a control. For individual units, average decoding performance was significantly better than chance (25%) and the control condition (average percent correct PC=29.5/28.8%, Wilcoxon signed-rank test performance shuffled vs. original Z=8.09/8.45 p<10^−6^, for high- and low-selectivity units, respectively). This was also true using population activity for stimulus selective units, while classification performance was not different from chance using population activity from less stimulus selective units (Wilcoxon rank-sum test Z=8.63/0.08, p<10^−6^/p>0.05, for high and low selectivity population, respectively). Similarly, the proportion of units showing significantly elevated decoding performance was higher amongst highly selective units (18.9% vs. 9%, CI95 15/24% and 6.3/11.5%, p<0.05 based on individual Permutation tests shuffled vs. original distribution). Comparing different MTL regions, we found that decoding performance was similarly high in hippocampal, enthorinal and parahippocampal units, but not different from chance for amygdala units (see **Fig. 4 f**). Taken together, our analyses suggest that phase of firing indeed differs depending on serial position of maintained memory items - as predicted by Lisman’s model.

### Phase-of-firing order does not correspond to item order

A second central prediction of the model by Lisman is that phase order matches the item order within the sequence. **Fig. 4 g-h** summarize our analyses addressing this question. We first show the phase distribution across neurons for each stimulus position based on theta frequencies exhibiting the largest power during the delay at the respective LFP channel (**Fig. 4 g**). Here, each neuron’s phase per position was normalized by subtracting the mean phase across all positions. Hence, if item position matched phase order, this would result in an equivalent ordering of phases across neurons. However, even though phase distributions were significantly different between positions as described for individual units above (circular ANOVA, Watson-Williams test F=2.89, p<0.05), our analyses did not reveal an item-position-equivalent phase ordering. Similarly, when analyzing phase order for individual neurons using mean phase of firing per position, we found that approximately 15% of units exhibited the stimulus-equivalent consecutive phase order (circular ordering clockwise, i.e. 1,2,3,4), 18.4% of unit-channel pairs showed the reverse order (i.e. 4,3,2,1) with both proportions not significantly different from the expected chance probability of 1/6 for a specific order (2 <0.1, p>0.05, **Fig. 4 h**). Thus, while Lisman’s model proposes an equivalence between phase- and stimulus order during memory, this was not reflected in our empirical results. Finally, we assessed the phase range used to encode item position in memory. **Fig. 4 h** shows the mean phase distributions across neurons where phases are sorted based on the preferred phase-of-firing order of each neuron instead of the actual item position. As expected from the spike-phase coupling we found, the phase range representing all positions spanned a fraction of the entire cycle of approximately 110 deg (IQ Range: 55.6 deg). Remarkably, within this restricted range, phase differences appeared to be equally distributed, with similar time lags between neighboring pairs of positions when mapped onto the time domain (Median time shift between pairs: 21.2ms, 20.4ms and 20.6ms, Kruskal-Wallis test Chi2<0.1, p>0.05, see **Fig. 4 j**).On the one hand, these findings support the notion that when spiking occurs preferentially at specific phases, it might be more efficient in impacting neural activity in target regions and foster effective communication and neural plasticity^25;26^. On the other hand, equal phase differences provide an efficient way to maximize the representation of information in phase space, as in our case 4 different item positions. Taken together, our analyses firstly reveal position-dependent phase-of-firing differences at the single-unit level during working memory. However, while our results support a phase-of-firing code for representing sequential items in memory as suggested by^1^, we find no equivalence regarding the ordering of spike-phase and position. This seems difficult to reconcile with the theory, which clearly predicts a correspondence between phase of firing and stimulus order.

### Phase modulation in a trained recurrent neural network model

Recurrent neural networks (RNNs) have previously been used to investigate neural computations during multiple cognitive tasks, including memory tasks^27–31^. Thus we set out using RNNs to assess potential neural mechanisms underlying our findings. We trained g rate-based RNNs on a task which was designed to be analogous to the one used during neural recordings (**Fig. 5 b-h** summarizes our training results, see also Methods and **Fig. S2 a** for details). In brief, four out of eight input units were sequentially activated during each trial, mimicking the presentation of stimuli in the experiment. This was followed by a delay and a presentation of the same four stimuli shown initially, in either a matching (‘correct’) or non-matching (‘incorrect’) order. During training the model was optimized to indicate matches and non-matches and training was run until 95% accuracy was reached on a validation set, containing stimulus combinations not used during the training epochs. We subsequently analyzed neural activity of the 40 trained networks in a similar fashion as our neural data: First, we identified units with increased firing rate during stimulus presentation that were also selectively responding to a specific stimulus. Over 40 models, on average 155 *±* 3.5 (mean *±* SE, N=40) out of 200 trained units exhibited stimulus-selective behavior (Wilcoxon signed-rank test, Baseline vs. Stimulus presentation time, p<0.001, for an example see **Fig. 5 c**). Next, we asked whether our models also exhibit oscillatory activity. We defined the model’s LFP as the summed absolute synaptic input to all neurons^32^ and observed oscillatory power within our trained networks at a range of frequencies typically peaking between 0.4 and 2 Hz, with an increase in power during delay as compared to baseline (Wilcoxon rank-sum test p<0.01; **Fig. S2 b**). In order to promote comparability between our oscillating networks and recorded data we further added a novel regularization term to manipulate the oscillation frequency, and chose several frequencies matching the observed theta frequency peaks in our neural recordings (see Methods for details).

**Figure 5.**
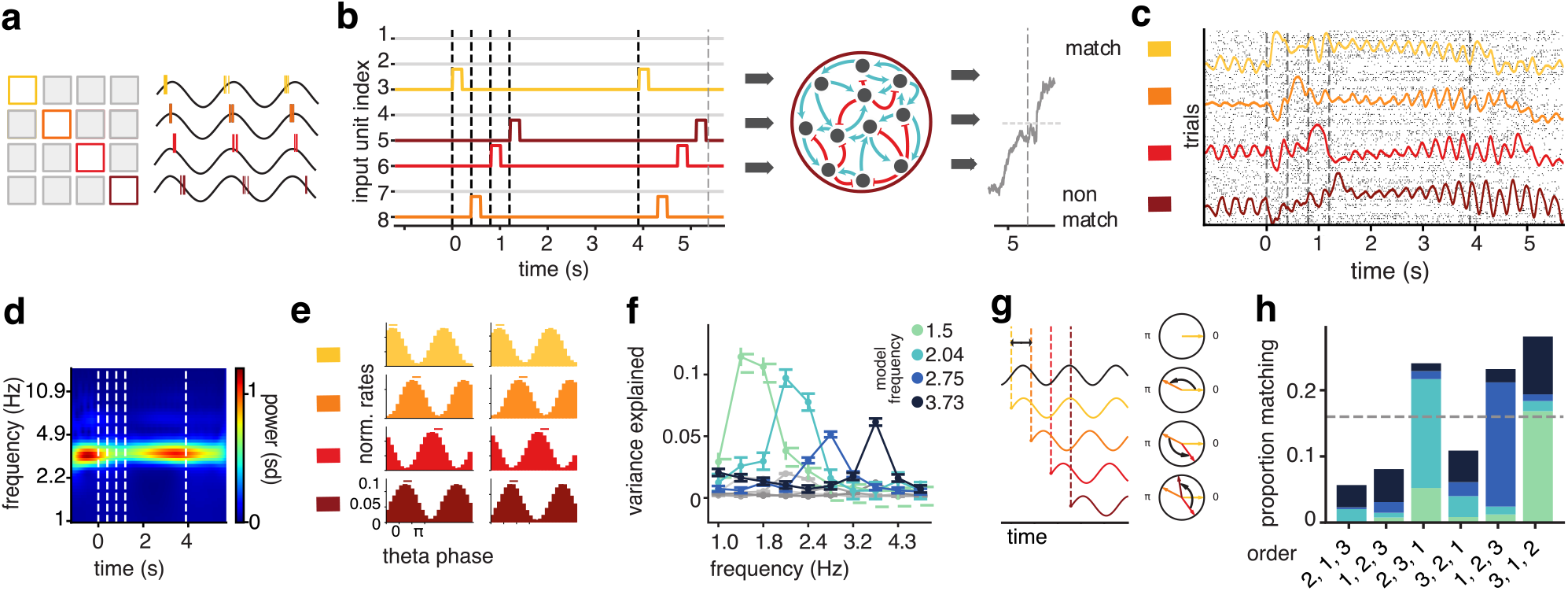
Phase-dependent firing in trained recurrent neural networks varies with item position. **a**. Schematic of the model proposed by Lisman & Idiart^1^. A stimulus-selective neuron spikes at different, consecutive theta phases in response to each of the 4 serial positions of a memory item (color-coded). **b**. RNN schematic showing example trial. Each colored line corresponds to an input unit activated with a stimulus, at different positions. **c**. Raster plots (spikes sampled from rate activity) and mean firing rate (colored lines) of a stimulus-selective model neuron shows a clear evoked response at each stimulus position, as well as oscillatory activity during delay. **d**. Time-frequency spectrum of trained RNN activity shows a power peak in the theta-range. **e**. Examples of phase-histograms of recurrent units during delay show different peaks of activity related to stimulus position. **f**. Circular variance explained (*V*_*ex*_) between stimulus positions is significantly elevated at tuned oscillation frequency (color-coded) in comparison to shuffled position labels. **g**. Schematic displaying proposed stimulus-induced phase reset and resulting phase order. A stimulus resets the phase of neurons to the same value, irrespective of the stimulus position within the sequence (colored lines), while the reference oscillation (black) is unaffected. As — depending on the oscillation frequency — the presentation of multiple stimuli given a specific inter-stimulus intervals (ISI) cannot always be contained within one oscillatory cycle, phase reset leads to a specific phase order as a function of the timing of the stimuli with respect to the reference oscillation. **h**. Proportion of matching position order based on phase of firing of RNN model showing a dependence on tuning frequency (x-axis) compatible with the prediction based on the relationship between stimulus onset differences and oscillation frequency, color coding as in **f**.

To analyze spike-phase coupling, we created spike-phase histograms by binning normalized firing rates of the upper 50th percentile of stimulus selective units during the delay, with respect to the phase of a sine wave with frequency matching the frequency with highest power in the models’ LFP spectra (**Fig. 5 d**). Similar to our experimental data, the firing rate of these units was coupled to oscillation phase and preferred phase of firing differed between sequence positions (**Fig. 5 e**). We quantified this effect by computing the circular variance explained *V*_*ex*_ between item positions for the upper 50th percentile of stimulus selective units. We typically observed a significant increase in *V*_*ex*_ as compared to shuffled position labels during the delay for models regularized in a range of different frequencies (**Fig. 5 f**). For the majority of these units (71.88%), phase order did not match item order within the sequence, similar to our neural recordings.

Our results demonstrate qualitative similarities between neural data and model activity and show that order-dependent (but not order-preserving) phases arise in a neural network trained to solve a sequential memory task. Can our models also help us understand how these non-ordered phase-relationships come about? We noticed in our models that phase of stimulus selective units was systematically reset by the onset of their preferred stimulus, and then remained at this stimulus-induced phase during the delay period (see also **Fig. 5 c**). In that case, the phase of such a unit, relative to an ongoing reference oscillation is determined by the timing of the stimulus with respect to the reference oscillation. When successive stimuli are shown, the phase differences between units coding for successive stimuli will then depend on the length of the inter-stimulus interval (ISI) relative to the cycle-duration of the ongoing reference oscillation (**Fig. 5 g, Fig. S2 c**). In line with this hypothesis, we observed that different phase orders emerged in our models as a function of oscillation frequency (**Fig. 5 h**; SE in **Fig. S2 d**). Finally, we also found this relationship in our empirical data: For each stimulus selective unit, we obtained the order of phase-of-firing at the theta frequency for which it exhibited the strongest phase differences between item positions (based on *V*_*ex*_). Subsequently, we quantified how many of these units exhibited a phase order as predicted by the frequency of the theta oscillation and the ISI used during the experiment (see **Fig. S2 c** for the predictions). For a significant proportion of units we were indeed able to predict the phase order correctly (25,2%, N=87, permutation test using shuffled labels between frequency and ordering, p<0.05, also see **Fig. S2 e**). Taken together, our analyses link the phase order to the ratio between cycle-duration of the oscillation and the ISI for both model and recorded neurons.

## Discussion

How is the temporal order of a sequence of items reflected in neural activity during a working memory task? In our study, we tested a long-standing theory^1^ on the role of spiking and theta oscillations using single unit and LFP recordings in human MTL recordings as well as recurrent neural network modeling. We observed that spike rates did not vary with item position during memory maintenance. Instead, neurons exhibited robust spike-phase coupling in theta frequencies, where preferred phase of firing differed between item positions but did not reflect item order. Importantly, this effect depended on whether the sequence of stimuli was correctly remembered or not, emphasizing the behavioral relevance of our findings. Many of our empirical results were supported by observations of RNN-based neural activity after training in an analogous task. This included strong stimulus selectivity during encoding, emerging oscillations, oscillatory phase coupling during memory maintenance and importantly, phase differences related to item position, where phase-of-firing order again did not match item order of the sequence.

Taken together, our findings corroborate one important prediction of Lisman’s model - namely that the serial position of memory items is encoded in phase of firing. Thus, in case of memorizing the order of multiple items, theta oscillations could indeed provide a temporal frame of reference for relating the ‘what’ *to* the ‘when’ as has been suggested previously^13^. Similarly, theta cycles could serve as a ‘separator’ between sequential memory items to facilitate read-out by downstream regions and thus serve both efficient encoding and routing of information within working memory^19;20;26;33;34^. However, in contrast to another prediction made by the Lisman model, our analyses did not show that the order of stimuli corresponds to phase-of-firing order for either recorded or modeled neurons. These observations might contradict two previous investigations on multi-item working memory^35;36^. Both studies used oscillatory activity in the gamma band in response to visual stimuli (50-100Hz, ‘gamma-bursts’) from iEEG / MEG signals respectively, to assess gamma phase ordering with respect to theta oscillations and observed corresponding item and phase order. However, neither study explicitly required subjects to memorize temporal order nor measured single unit activity. The latter is particularly important, since linking spiking of individual neurons to gamma band activity is challenging and depends on many factors including intrinsic neural network properties^37–39^. Specifically, whether gamma bursts indeed reflect select responses to different visual stimuli, such as those we have demonstrated for spiking of individual neurons in MTL, remains unresolved. Our findings might provoke the question of why sequence position would be represented by phase-of-firing if the order of phase does not correspond to stimulus order? It has been argued MTL activity is not stringently dependent on the concept of physical space or time but rather represents events relative to each other, such as in our case stimuli at their respective position^13^. This scheme still applies to our findings. That is, if there is an arbitrary—yet consistent—spike-phase relationship per neuron between different positions, then relational coding, as in reading out position of stimulus A (by neuron X responding to stimulus A) vs. position of stimulus B (as in neuron Y encoding stimulus B) would still be possible at the population level.

Indeed, our oscillating RNNs could maintain sequential information in working memory while having ‘non-ordered’ phases of firing. Moreover, we derived and tested a hypothesis on how phase order might arise based on model observations, providing a possible mechanistic explanation for our results. Stimulus-induced phase reset in MTL neurons has been reported previously, in particular in association with increased theta synchronization^40–46^. In comparison to previous work which had demonstrated maintenance of mnemonic information in point, line and plane attractors^47–50^, we thus show emergent oscillatory dynamics during working memory (in line with recent work^51^) and provide a novel and alternative coding mechanism within artificial neural networks. An interesting question for future studies will be to investigate whether and how different dynamical coding schemes are related to each other, under which experimental conditions and manipulations either one arises and how they relate to different neural mechanisms during memory processing. Our modeling framework makes it possible to further study oscillatory network dynamics at a mechanistic level during memory processes and emphasize the general importance of RNN-based modeling in systems-level neuroscience research.

Ultimately, for both *recorded* and *modeled* neural activity, our observations indicate a temporal code for non-spatial sequential information within memory in analogy to previous reports on spike-phase coding of place fields during spatial navigation and memory in rodents^13^. Thus, our observations point to a more general role of temporal coding based on oscillations within the medial temporal lobe.

## Acknowledgements

This work was supported by the German Research Foundation (DFG): SFB 1089, SPP2205, MO 930/4-2, the Volkswagen Foundation: 86 507, and the German Federal Ministry of Education and Research (BMBF): 031L0197B, Tübingen AI Center, FKZ: 01IS18039A. SL, MP and JHM are associated with the Machine Learning Cluster of Excellence, EXC number 2064/1 – Project number 390727645. MP is supported by the International Max Planck Research School for Intelligent Systems (IMPRS-IS). We thank Richard Gao and Janne Lappalainen for discussions and comments on the manuscript.

## Materials and Methods

### Participants

16 patients (9 female, 7 male, median age = 42/45 years, respectively) with chronic, intractable epilepsy were implanted with depth electrodes to undergo seizure monitoring for presurgical evaluation. All subjects gave their written, informed consent to participate in the experiments. Subjects performed either 224 trials or 112 trials (one subject) of a modified Sternberg task as described below.

### Ethics statement

The study was approved by the Medical Institutional Review Board of the University of Bonn (accession number 095/10 for single-unit recordings in humans in general and 249/11 for the current paradigm in particular) and adhered to the guidelines of the Declaration of Helsinki.

### Paradigm

The paradigm was a multi-item sequential memory paradigm (modified Sternberg paradigm) as illustrated in **Fig. 1 a**. On each trial, a fixation cross appeared for 1000 ms. This was followed by a temporal sequence of four different stimuli that were chosen randomly on every trial out of a set of eight stimuli in total. Pictures were chosen based on a pre-screening run one to four hours before this experiment (see^52^). Pictures contained mostly natural images depicting photographs of people, places or objects. Within the sequence each stimulus was presented for 200 ms with an inter-stimulus interval of 200 ms. The presentation of the forth stimulus was followed by a delay period showing a blank black screen. The delay period ranged between 2400 and 2600 ms (Median=2500 ms, IQR = 100 ms). Finally, a stimulus panel simultaneously showing four possible stimulus sequences was shown on the screen, one of which matched the one previously shown (Chance level = 0.25). Subjects were instructed to press a key number indicating the row of the matching sequence. Each of the 8 different stimuli used was presented equally often at each of the four temporal positions. On each trial the sequence was randomized, such that subjects did not know the upcoming sequence on any given trial. In total patients completed 224 (112 for one subject) trials. Each stimulus was shown within the sequence on half of the trials (112 out of 224 trials) and not part of the the sequence of images on the other half of the trials. Thus, we showed 112 (56 for one subject) trial unique sequences from a possible set of 1680 (8!/4!) within each experiment, where each image was shown an equal number of times at each position within the sequence (N=28 trials * 4 positions). The matching sequence in the probe display was shown counterbalanced in each row of the panel (i.e. each row approximately 25 percent). Within the probe sequences, images were counterbalanced in a way that the task could not successfully be solved (i.e. max. avg. 50 % correct) when simply remembering the first, the last or the first two stimuli within the sequence.

### Recording technique

Recording techniques presented here have been described in detail in previous studies (see for example^53^). In brief, we recorded the raw voltage traces from nine microwires (8 high-impedance recording electrodes, 1 low-impedance reference; AdTech, Racine, WI) protruding from the shaft of depth electrodes recorded with a sampling rate of 32 kHz. Signals were amplified and recorded using a Neuralynx ATLAS system (Bozeman, MT) and referenced against one of the low-impedance reference electrodes.

#### Data Analysis

All data analysis was performed using custom-written functions as well as the CircStatsToolbox written for Matlab Version R2014b as well as pycircstats - Toolbox written for Python^54^.

#### Behavioral analysis

For each trial we obtained the keyboard response (i.e. sequence number 1-4) of the subjects and quantified the proportion of correct responses (PC) as the number of trials for which the subjects’ response matched the previously shown sequence. Reaction times were computed as difference in time between probe onset and subjects’ key press.

#### Spike analyses

Throughout the manuscript, we use the terms ‘neuron’, ‘unit’ and ‘cell’ equivalently to describe recorded responses of presumed neuronal spiking. We use spiking equivalently to firing, in order to describe neuronal activity of single neurons. Model units describe the output activity of the artificial RNN units. Whenever multiple comparisons were performed, p values were corrected using the Simes procedure^55^.

Spike sorting was performed semi-manually using Waveclus 2.0 and Combinato^56;57^. Based on thorough manual visual inspection of waveforms we removed unit recordings that were contaminated by artefacts or were temporally unstable over the course of the recording time, which resulted in a selection of 1420 units from 921 unique LFP channels in medial temporal lobe regions (hippocampus **HPC**, N=564/376, 40%, enthorinal cortex **EC**, N=251/161, 18%, parahippocampal cortex **PHC**, N=213/138, 15% and amygdala **A**, N=392/246, 28%, units / channels, respectively).

#### Identification of responsive and selective units

In order to assess whether a unit significantly responded to a stimulus, we compared spiking activity during pre-stimulus baseline to the stimulus period, where baseline activity was defined as the average spiking activity within a 500 ms window prior to the onset of the first stimulus and stimulus related activity was defined as activity following a stimulus. Spikes were binned into windows using a bin size of 100ms with 50ms overlap starting at 100ms until 800ms poststimulus onset. Using a one-tailed Wilcoxon signed-rank test we compared median spike rate within each bin vs. baseline across all trials in which a particular stimulus was shown irrespective of stimulus position (N=112*4). Associated P-values were computed for each window and corrected for multiple comparisons using the Simes procedure^55^, N=13 time windows). A unit was defined as responsive if there was a significant increase in spike rate relative to baseline (p<0.001) for at least one window within the range of 250- 600ms post stimulus onset for hippocampal, entorhinal and amygdala units or 170-550ms post-stimulus onset for parahippocampal units, taking known ^**? ?**^ and observed regional differences in stimulus response latency into account. This procedure resulted in N=217 highly stimulus-responsive units (N=82 hippocampal units, N=55 amygdala units, N=55 parahippocampal units, N=25 entorhinal units).

#### Latency of stimulus evoked response

To assess stimulus response latencies, we computed average spike rate (Hz) in consecutive 20 ms windows with 10 ms overlap and compared the activity in each time window with baseline activity as decribed above. The latency of the response was then defined as the time bin for which there had been three consecutive prior bins with an associated significant increase (p<0.001) in spiking activity. The resulting median latencies were 222 ms for hippocampal units, 258 ms for entorhinal units, 190 ms for parahippocampal units and 240 ms for amygdala units, (IQRs: H:128 ms, EC:171 ms, PHC:90 ms, A:141 ms).

#### Stimulus selectivity

For each unit, we defined the stimulus eliciting the largest firing rate increase relative to baseline (based on the minimal p-value) as the ‘preferred stimulus’ (**PS**). Since units sometimes responded to multiple stimuli, we also assessed how selective their responses were to a particular stimulus out of all 8 stimuli shown. We computed the normalized difference in spiking (Hedges’ g, equivalent to Z-score scale,^58^ by first subtracting the mean evoked spiking activity to the stimulus eliciting the second largest response from activity to the PS and dividing by the pooled standard deviation across trials containing both stimuli. Window sizes used for obtaining stimulus-related activity were the same as described above. Based on this selectivity metric, we grouped units into a “high” vs. “low” selectivity group based on whether individual selectivity indices were larger or smaller than the 50th percentile across all units (median split). By comparing activity during trials in which the PS was shown within the sequence (half of the trials) to trials in which it was not, we were able to track stimulus-specific effects in neural activity during the maintenance period. Using the selectivity metric further allowed us to test stimulus specificity for more or less selective neurons. For the unit shown in **Fig. 1 a** the associated Hedges’ g was 1.01, indicating that the response to the preferred stimulus was about 1SD larger than to the stimulus with the second highest response. The selectivity index of this unit fell within the upper 25th percentile of the distribution across the population (see also **Fig. S1 d**).

To estimate single unit activity across the trial period, we binned spikes with a resolution of 1 ms and obtained instantaneous firing rates by convolving the spike trains with an Gaussian kernel (SD = 25 ms) per trial. We transformed instantaneous firing rates into Z-Scores by normalizing to the mean activity and standard deviation during the 500ms baseline period. To compare spiking between different trial windows, Z-Scores were averaged during the visual response window (see above), the delay period (1500ms window prior to probe onset) and after the probe onset (200-500ms).

#### LFP analysis and spike phase coupling

Spectral analyses were similar to a previous report^33^. In brief, we obtained the time-frequency decomposition of the downsampled (1000 Hz) LFP signal using complex Morlet wavelets (c=7 wavelet oscillations) and extracted the instantaneous amplitude and analytical phase as a function of time and frequency by convolving the raw real-valued time series x(t) with the complex Morlet wavelet w(t, f0) to obtain the complex output signal y(t, f0), also denoted as the analytic signal, where f0 denotes the desired center frequency of the wavelet function. The center frequencies f0 to obtain power spectra as shown in **Fig. 2** were created by exponential spacing of 100 frequencies between f=2^x^ with x = 6/8,…,54/8 resulting in a range of frequencies approximately between 1.5 and 108 Hz. Oscillatory power change during the delay was assessed by transforming the raw power spectra to Z-score scale relative to baseline power by subtracting the mean power during baseline (500 ms window preceding the first stimulus) and dividing by the standard deviation during baseline across trials from each individual trial before averaging across all trials (see **Fig. 2a**). Delay spectra as shown in **Fig. 2b** were obtained by averaging across 1500 ms prior to probe onset (equal to spiking activity window). For the spike-phase analyses, we specifically focused on theta frequencies. As the exact frequency range associated with ‘theta’ differs among studies, and different frequencies within the theta range might even be associated with inter-species differences^59^ we obtained the phase of the analytic signal over a wide range of frequencies starting with 1.5, 1.75, 2.03, 2.37, 2.8, 3.2, 3.7, 4.4, 5.1, 5.9, 6.9 and 8 Hz. We analyzed simultaneously recorded spiking and LFP at the same microwire from 217 channel-unit pairs, where 185 LFP recordings were paired with one unit recorded at the same electrode and 32 LFP recordings were paired with more than 1 unit (27 channels with 2 unit pairings, 5 channels with 3 unit pairings). To achieve higher statistical robustness from larger spike counts and to be able to estimate spiking at lower theta frequencies, we subsequently selected units for which average firing rates were above 2 Hz within the delay period (85%) and used a baseline period of 1000ms and a delay period of 2000ms for comparisons (choosing the last 2s before panel onset and leaving a minimum of 500ms interval *post*-stimulus *offset*). For each of the units we obtained spike phase histograms throughout the time periods by binning spiking at theta phase into equally spaced phase bins with a width of pi/16. To account for non-uniformity of phases not associated with spiking, average spike-phase counts were normalized per bin by the number of occurrence of each particular phase bin during the considered trial window. We subsequently quantified the preferred phase angle (mean directionμ) as well as the magnitude of spike phase coupling (concentration parameter *κ*) by fitting von Mises density functions to the spike distribution across phase bins. This procedure was repeated across all trials during which the PS was shown vs. not (NPS trials) and separately for trials with different stimulus positions of the PS within the sequence (1-4). We applied the Rayleigh test for circular uniformity to the distribution of the mean preferred phase angles across all neurons within each region and used Rayleigh’s Z score as a test statistic to compare uniformity of spiking between different regions. To compare mean preferred phase angle between regions, we estimated the population phase angle by averaging across mean phases of all neurons within each region.

Comparisons of spike-phase-coupling magnitude (i.e. *κ*) can be confounded by differences in spike rates between compared conditions. Indeed, we observed small, yet significant differences in spike rates between baseline and delay activity across all units (median spike rate in Hz baseline / delay 2.8/3.6, IQR 4.7 / 5.5, Wilcoxon signed-rank test Z-Value 8.06, p<10^−5^) as well as within different regions (Median spike rate in Hz Baseline/Delay HPC: 4.25/4.67, EC: 1.81/1.98, PHC: 2.82/3.32, AM: 2.60/2.83, IQR HPC: 7.46/7.13, EC: 3.59/5.49, PHC: 4.46/5.47, AM: 3.24/3.84; all units Wilcoxon signed-rank test Z=8.09, p<10^−15^, for individual regions: HPC:Z=4.6152, EC:Z=2.29, PHC:Z=4.3726, AM:Z=4.3401, all p<0.02). To assess whether differences in spike-phase coupling could have resulted from the observed spike rate differences, we performed the following control analyses: We first computed the average spike rate (in Hz) for baseline and delay (see above) and subsequently split units into the upper and lower 50th percentile based on spike rate differences. As expected, for units in the lower 50% group, we found no significant difference in spike rates between windows (‘Glptile’, N=36/11/33/27 for HPC, EC, PHC and AM, p>0.05 based on Wilcoxon signed-rank test). Comparing median *κ* values between baseline and delay for this group (‘Glptile’), we still observed significantly increased spike-phase coupling during delay (Z>5.15, p<10^−6^). In a second control analysis, we ranked units based on their difference in spiking activity between both windows and consecutively excluded units until we no longer observed a significant difference in spike rate (‘Gexcl’, N=51/20/48/39 for different regions, Wilcoxon signed-rank test p>0.05). Again for this group (‘Gexcl’) we observed significantly larger median *κ* during delay compared to baseline (Wilcoxon signed-rank test, Z>8.69, p<10^−16^). We performed the same analyses controlling for differences in delay-related spiking between PS vs. NPS trials (Wilcoxon signed-rank test based on median spike rates during delay: N=217, Z=5.66, p<10^−7^) and still observed a significant increase in spike-phase coupling between these two conditions (Wilcoxon signed-rank comparison median *κ* in ‘Glptile’ N=108, Z=6.55, ‘Gexcl’ N=7.89, p<10^−10^). Finally, we also tested median *κ* values between the low vs. high selectivity groups of units after eliminating spike rate differences using the same procedure as described above (Median spike rate low vs. high, 4.52/3.6 Hz (qqq fm: isn’t this high vs. low?), Wilcoxon rank-sum test Z=-2.77, p<0.01). While we initially observed that high-selectivity neurons exhibit slight albeit significantly stronger spike phase coupling than the low-selectivity group (Wilcoxon rank-sum test comparing median *κ* Values Z=2.13, p<0.04), this effect was abolished after correcting for spike rate differences between the two groups (Wilcoxon rank-sum p>0.05, Z=1.86).

#### Analysis of phase of spiking at different stimulus positions

To analyze whether preferred phase of firing during the delay differed from stimulus positions during encoding, we obtained trial-based estimates of the mean preferred phase of spiking during the delay for each neuron during trials showing the preferred stimulus of that unit within the sequence. We subsequently computed the circular variance explained between different experimental conditions (in our case stimulus positions 1-4) by quantifying the ratio of variance within conditions relative to the variance across conditions. We defined circular variance within condition, i.e. position, by 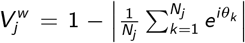 where *N*_*j*_ is the number of trials in condition *j*, the index k runs across all trials of condition *j*, and *θ*_*k*_ is the mean phase of spiking in trial *k*. We subsequently calculated the mean variance within conditions as 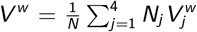 where *N* is the total number of trials and *N*_*j*_ the number of trials within condition *j*. While there were typically 28 trials in each condition, an unequal number of trials could potentially arise from the fact that trials during which no spikes were detected did not contribute to the mean phase estimate (on average 10% of trials across neurons). We further defined circular variance across all conditions by 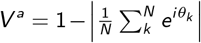 where *k* runs across all *N* trials. We finally computed the circular variance explained per neuron as follows: 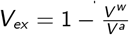.

For non-parametric statistical comparisons we also obtained a random distribution of *V*_*ex*_ by shuffling trial labels N=1999 times between different conditions based on random permutations. We repeated this procedure for every unit using all trials, and separately for correct and incorrect trials (including the respective shuffle-based random distribution to account for trial count differences between these two conditions). For each permutation run, we also obtained an estimate for the proportion of units showing a significantly larger *V*_*ex*_ compared to their shuffled distribution. A unit was defined as significant if the true *V*_*ex*_ exceeded the shuffled *V*_*ex*_ estimates in at least 95% of the cases (p<0.05) for at least two frequencies. To obtain confidence intervals on proportion estimates, we created surrogate distributions of units using a bootstrap procedure (with replacement, as implemented by *bootstrp* in Matlab 2014b, N=1999) comparing the units’ *V*_*ex*_ and respective shuffled distributions for each drawn sample (see Supplementary **Fig. S1e**).

#### Phase order estimation

As described above, we obtained the mean phase per neuron for each stimulus position of the PS during the delay. To estimate phase order, we first subtracted the mean phase across positions from each position’s phase, thereby anchoring all phases relative to 0 deg phase (see also **Fig. 4g**). We subsequently sorted phases counter-clockwise in an ascending order and obtained the stimulus position index for the sorted phases accordingly. A phase order was defined as equivalent to position order if the relative ordering along the circle matched the stimulus order (expected by chance in 1/6 of cases), and reverse if the reverse stimulus order was matched (likewise expected in 1/6 cases).

#### Decoding analysis

To test whether stimulus position can also be decoded from phase of spiking, we used a support vector machine (SVM) algorithm as implemented in Matlab 2014b. Here, phase of spiking (mean phase per trial per neuron at stimulus position of the individual units’ PS) served as the predictor matrix to predict one of four possible binary class labels (i.e. stimulus position) in a multi-class classification (chance performance 25%). The learning algorithm used a rbf kernel and a one vs. one encoding scheme for the predictor matrix. We randomly partitioned the data into a training and test set for cross validation using a holdout proportion of 0.15 (training on 85% of the data, testing on 15%) N=101 times. Decoding performance was defined as the proportion of correctly assigned class labels (i.e. position indices) on the test data set after training. To perform statistical comparisons, we repeated the decoding procedure after shuffling true class labels across trials, i.e. randomly assigning position indices to trial phases in order to create a random distribution of prediction performance. We obtained the associated p value for the comparison between true vs. shuffled labels by the number of cases for which the true performance was lower than the shuffled performance. For population-based decoding, we created a pseudo-population of units by pooling all units across all sessions and subjects and taking the phase of spiking at all theta frequencies (see above) into account. Decoding performance was separately assessed for groups of units with high vs. low stimulus selectivity, and also per brain region. In addition, we quantified decoding performance of single units using the same approach and compared median performance between true vs. shuffled label runs across all units, separately for the high vs. low selectivity group. To estimate the effect size of position decoding between regions, we calculated the difference between true performance in standard deviation units (Hedges g, as described above) and obtained confidence intervals based on bootstrapping with N=1999 repetitions.

## Recurrent Network Model

### Code availability

All code for training and analyzing the RNN models is available at https://github.com/mackelab/sequence-memory.

### Model definition

Our models consists of a recurrent network of *N* firing-rate units:

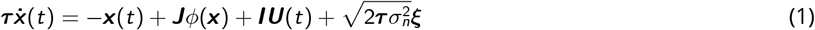

where ***τ*** ∈ ℝ^*N*^ is a vector of time constants, ***x*** ∈ ℝ^*N*^ denotes the current of each unit, *ϕ* is the (non-linear) activation function and ***J*** ∈ ℝ^*N*×*N*^ is the recurrent weight matrix specifying the connectivity between units in the network, ***I*** ∈ 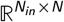 the input weight matrix specifying the connectivity from stimulus input to recurrent units, and 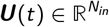 the time-varying stimulus input. ***ξ*** denotes *N* Gaussian noise processes with zero mean and unit variance, representing intrinsic network noise scaled by 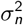. An overview of all parameters can be seen in Table S1.

We implemented biophysical constraints on the weight matrix ***J*** in line with previous work^30^: We allowed neurons to have either only excitatory or only inhibitory outgoing connections (Dale’s law). To achieve this, we initialized a matrix ***J***_***opt***_ with samples drawn from a half-normal distribution *Y* = |X|, such that *X* is a zero-centered normal distribution with variance 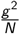. The matrix ***J*** is then computed as the dot product of ***J*** _***opt***_ with a diagonal matrix ***D*** where the first *N*(1 − *p*_*inh*_) elements in ***D*** were set to a positive number, and the last *Np*_*inh*_ elements were set to a negative number. Here *p*_*inh*_ denotes the fraction of inhibitory neurons. We choose elements in ***D*** such that the expectation of the recurrent input to neurons stays 0^30;60^, and the average of the variance of the excitatory and inhibitory populations is 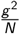. Mathematically, this results in the bulk of the eigenspectrum of ***J*** lieing in a circle on the complex plane with radius *g*^60^. The resulting dynamics are then given by a sharp transition from stable to chaotic dynamics at *g* = 1 as *N* → ∞^61^. For finite networks (as used in our study), instead of a sharp transition, there is an intermediate region where one finds limit cycles. During training, we optimize ***J***_***opt***_, and compute ***J*** as |***J***_***opt***_ |_+_***D***; we rectify the elements in the ***J***_***opt***_ by applying the ReLU function to ensure that the excitatory-inhibitory constraint is fulfilled.

### Task

We adapted the task performed by the human participants such that it could be readily performed by an RNN, keeping the information that had to be maintained during the delay period (four stimuli and their order) identical. The input to the network at a particular time step, ***U***(*t*), was always 0, except during the stimuli and probe periods. During these periods four out of eight stimuli were activated sequentially for 0.2 s by setting the corresponding entry in the vector ***U***(*t*) to 1. The activated stimuli were randomly chosen (without replacement) with equal probability every trial. After a delay period, the same four stimuli as during the stimulus period were shown, but now either in a different order (randomly drawn) or in the same order, with equal probability.

### Training procedure

During training we simulated equation 1, using the Euler method with step size △*T*. Giving us at time step *t*

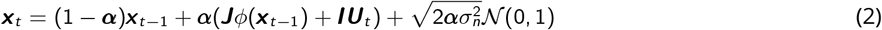

With 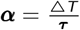.The network’s output is described by a linear readout of the firing rates *y*_*t*_ = ***W*** *ϕ*(***x*** _*t*_) with ***W*** ∈ ℝ^1×*N*^.

To keep model time constants within a biologically plausible range [*τ*_*min*_, *τ*_*max*_], we applied a nonlinear projection map^62^. We chose ***τ*** to be

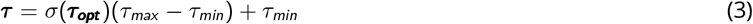

We initialized ***τ***_***opt***_ with samples drawn from 𝒩(0, 1) and optimized this during training. With *σ*being the logistic function, *τ* approaches *τ*_*max*_ as *τ*_*opt*_ → ∞ and *τ*_*min*_ as *τ*_*opt*_ → −∞.

In order for the network to perform the task, it had to determine whether the order of the initial sequence of stimuli matched the sequence presented after the delay period. We defined a scalar *ŷ*_*t*_ that was either 1 (match) or −1 (non-match) during the decision period.

We defined the loss of a single trial as follows:

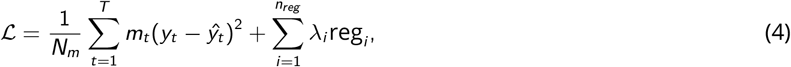

where *m*_*t*_ is 1 during the decision period, otherwise 0, and *N*_*m*_ denotes the number of non-zero *m*_*t*_. reg_*i*_ denotes additional regularization terms applied with weight *λ*_*i*_. We applied an L2 penalty on the rates to prevent implausible saturation of the activation function 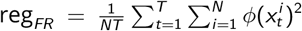 (^29^) and additional regularization to control the model’s oscillation frequency (equation 6).

We optimized parameters of the network by minimizing equation 4 using gradient descent. We calculated the average gradient at a particular time step over a subset (batch) of trials using back-propagation through time (BPTT), with the Adam^63^ optimizer in tensor flow^64^ with default settings (first and second order moment equal to 0.9 and 0.999, respectively).

Training with BPTT notoriously suffers from exploding and decaying gradient^65^. In order to avoid the former we used gradient clipping. If the norm of the gradient 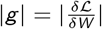 was larger than some maximum *g*_*max*_, we multiplied *g* with *g*_*max*_ */*|*g*|. To avoid vanishing gradients we employed curriculum learning and first train on a short delay (0.2s). Sweeps over multiple models were realized used the Weights & Biases toolbox^66^.

### Computing local field potentials in RNNs

Local field potentials (LFPs) as recorded from the brain are generated from currents of neurons embedded in a three-dimensional space, where the exact arrangement of neurons hugely influences the recorded signal. Neurons in our model, however, are completely agnostic to physical space.^32^ compared various LFP proxies for standard leaky-integrate-and-fire (LIF) networks, and found that a specific linear combination of the LIF synaptic currents provides an accurate LFP proxy. However, the authors also suggested a simpler proxy that is plainly the (absolute) summed AMPA and GABA currents. Given that our units have even less detail then the LIF neurons, we calculated the LFP in line with this, by taking the summed absolute synaptic input as LFP:

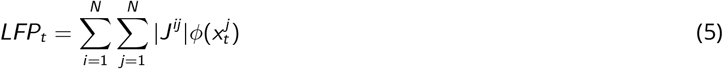

To systematically analyze the effect of oscillation frequency on the representations used by our model we developed a new loss term to promote a LFP with a peak at a specified frequency during training. This allows one to shift the natural oscillation frequency of the model to a specific frequency. We applied this regularization to all models described in the main text.

For every training iteration we computed the LFP. To make the loss term amplitude invariant we first normalized the LFP: 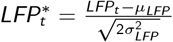. We then took the norm of the Fourier component at a specified frequency:

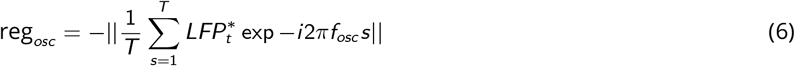

where *s* is the time associated with time step *t* in seconds, *T* is the trial duration, and *f*_*osc*_ the regularisation frequency in Hz. We selected frequencies congruent with the range of frequencies found in our experimental data, and trained 10 models for each frequency (1.5, 2.04, 2.75, 3.73 Hz), 70% (28/40) of which exhibited peak oscillations at target frequency after training (mean delay LFP power at the target frequency, pre training: 0.13 *±* 0.022, post training: 0.68 ± 0.048, mean ± SE, N=28).

### Model analysis

Analysis of the model was performed in Python using the pycircstat package^67^ as well as custom-written code. For the subsequent analysis we first generated 224 unique stimulus combinations matching the number of trials in the experiment. Half were randomly assigned to be match trials (the later four stimuli are presented in the same order as the initial stimuli) and the other half were assigned to be non-match trials (the order in which the initial and post-delay stimuli were presented differed). We repeated the analysis performed on the experimental data with the following adaptions to account for the model having a continuous firing rate instead of discrete spikes: To find stimulus-selective neurons, we used mean firing rates and counted activity starting with stimulus onset as stimulus-triggered activity. To create analogues to spike phase histograms, we binned continuous firing rate with respect to the phase of a reference oscillation, making sure that our bin size never exceeded our simulation time step. The reference consisted of a sine wave with frequency corresponding to highest power in the model’s LFP spectrum. Units in our model used tanh, with range (−1,1) as activation function, due to desirable properties with regards to propagation of gradients during training. During analysis the firing rates where first mapped to (0,1), using a linear projection map.

## Author contributions

F.M. conceived the study and administered the project. J.N. and F.M. designed the experiment. J.N. implemented the experiment. J.B. and F.M. performed the surgeries. F.M. and C.E.E. recruited patients. J.N., J.F., T.P.R. and F.M. collected data. M.P. designed and implemented the RNN model, carried out analyses on RNN model units and wrote the modeling section. S.L. and J.H.M. supervised modeling. S.L. and J.N. curated data. S.L. and T.P.R. performed the decoding analysis. S.L. conceived, designed, implemented and performed formal analyses and visualization, and wrote the paper. J.H.M and F.M. provided critical review and editing of the manuscript draft. All authors reviewed the results and approved the final version of the manuscript.

## Supplementary Information

**Table S1.** Parameters used to train the RNNs

| Key | Value |
| --- | --- |
| $N$ | 200 |
| $N_{in}$ | 8 |
| $\phi()$ | $\tanh()$ |
| $g$ | 1.5 |
| $p_{inh}$ | 0.2 |
| $\sigma_n$ | 0.05 |
| $\tau_{min}$ | 20 |
| $\tau_{max}$ | 120 |
| optimize | $I, J_{opt}, W, \tau_{opt}$ |
| batch size | 128 |
| learning rate | $5 \exp -5$ |
| $\lambda_{FR}$ | $1 \exp -5$ |
| $\lambda_{LFP}$ | $1 \exp -1$ |

**Figure S1.**
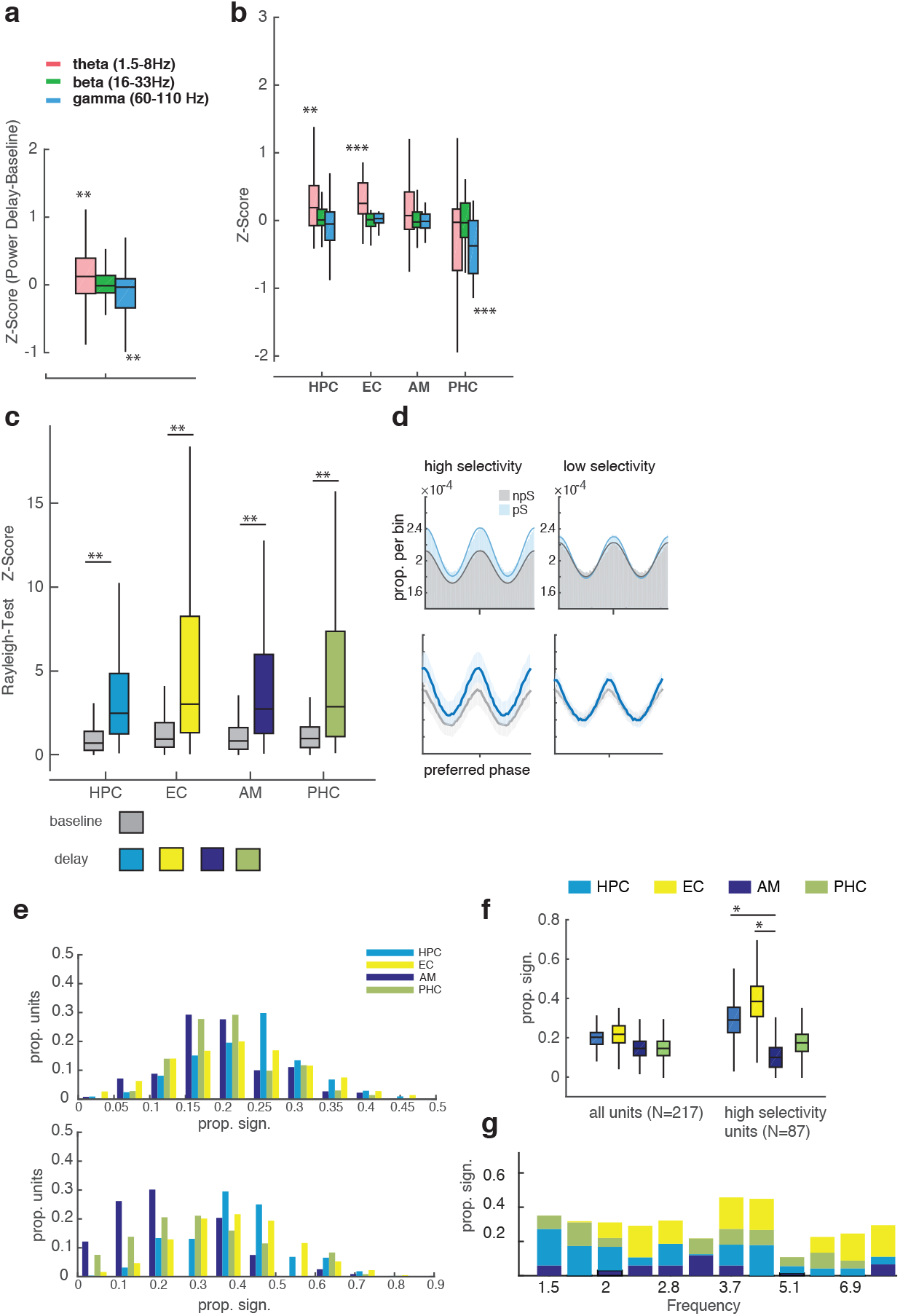
**a**. Normalized difference in oscillatory LFP power between baseline and delay (Z-score), showing significantly increased power in the theta band (including lower frequencies) and significant reduction in the gamma frequency band (60-110 Hz) during the delay (Wilcoxon Signed Rank Test, p<0.01, N=185 unique LFP channels, based on visually responsive units) **b**. Same plot as in **a**. split by region. Mainly, significant theta power increases can be found HPC and EC across all channels (Wilcoxon signed-rank test p<10^−5^/p<0.001, respectively) whereas amygdala and parahippocampal cortex showed increases as well as decreases in theta power (p>0.05). Interestingly, sign. gamma decreases during the delay can be found in PHC. **c**. Non-uniformity of theta-related phase of firing as indicated by Rayleigh’s Z-Score separately estimated from baseline and delay activity per region. In all MTL regions investigated, we find significantly enhanced spike phase coupling during delay compared to baseline (N, stats, all trials used, irrespective of PS/NPS). **d**. Spike phase histograms (upper panels) and van Mises fits (lower panels) estimated from theta band spiking activity during delay separately plotted for trials containing the PS vs. not during the stimulus sequence and neurons with high (blue) vs. low (gray) stimulus selectivity. Shaded areas correspond to SEM across units. Although the spike modulation seems to be larger for the high selectivity group of units, estimates based on individual concentration parameters *κ* were not significantly different after correcting for spike rate differences. **e**. Histograms showing the distribution of proportions of units showing significant differences in phase of firing between stimulus positions based on *V*_*ex*_ - Permutation tests per region. Histograms are shown for all units (upper panel) and again for units exhibiting high stimulus selectivity (lower panel). **f**. Proportion of units shown per region showing significantly elevated *V*_*ex*_ when compared to shuffled trials based on Permutation tests (criterion p<0.01). Among highly stimulus selective units, the highest proportions were found in hippocampus and entorhinal cortex. **g**. Same data as in **f**. for stimulus-selective units, per theta frequency and region.

**Figure S2.**
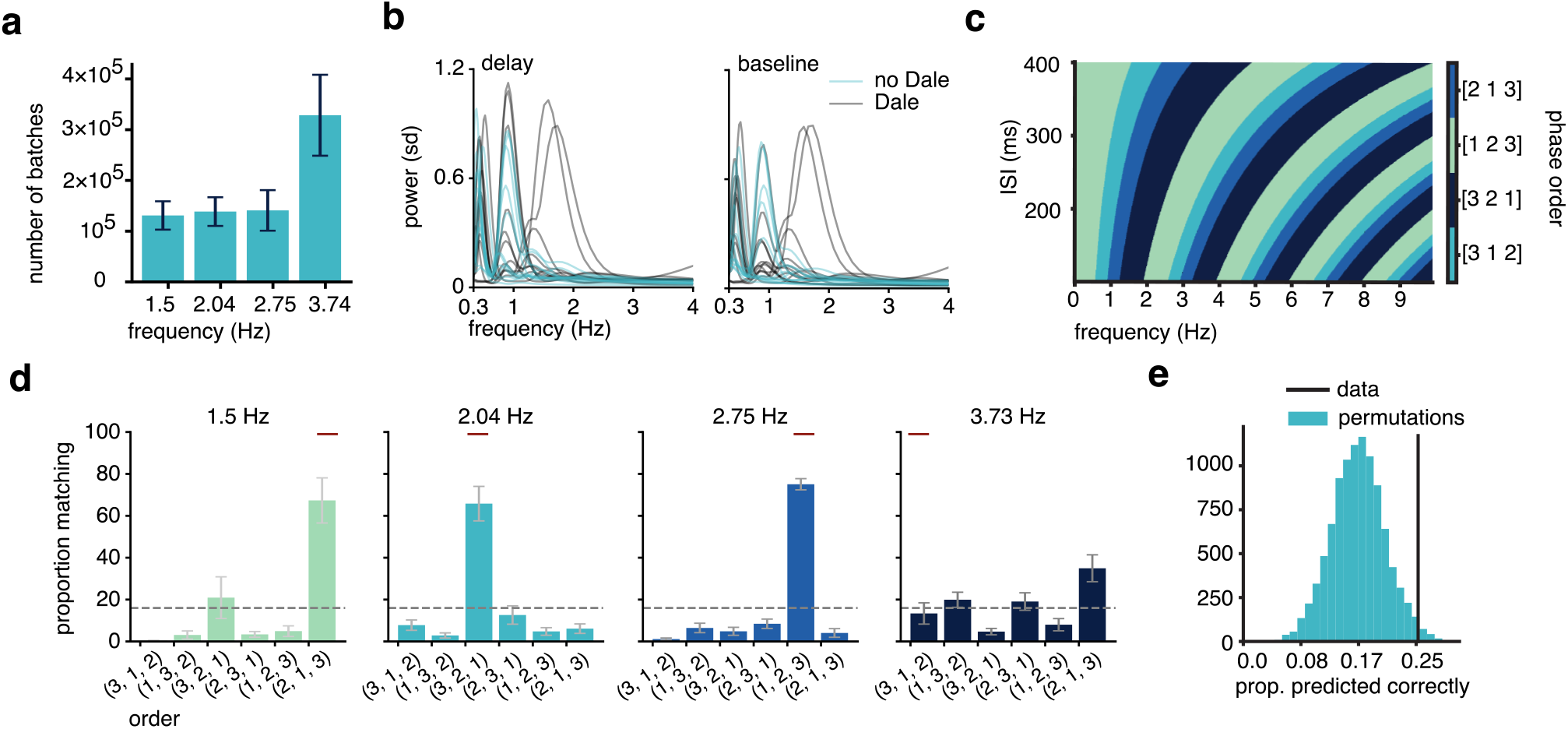
**a**. Mean (+ SE) training duration of models per regularization frequency. We noticed a frequency dependent effect of training on convergence rate in line with existing work^68^. **b**. Mean power during baseline and delay for 24 models demonstrate trained models oscillate, when trained without oscillatory regularization (12 with and 12 without Dale’s law, *g* =1, other parameters as in table S1). The increase in power during delay when compared to baseline is significant (Wilcoxon rank-sum test with N=24, p<0.01). To accurately detect the lower frequencies we first extended the baseline and delay period to 10s before calculating power. **c**. The predicted phase order depending on oscillation frequency and inter-stimulus interval, based on the simplified model (**Fig. 5 g**. Boundaries between the different phase orders at solutions of *exp*(*ip*2*πfs*_*isi*_) = 1, where *p* ∈ {1, 2, 3} denotes stimulus position, *f* is oscillation frequency at stimulus onset in *Hz* and *s*_*isi*_ is the interval between successive stimulus onsets in *s* **d**. Mean (+SE) proportion of phase order exhibited by models per regularization frequency. Which phase order occurred most frequently for the three lower frequencies is in line with the prediction of our simplified model (red bars). **e**. Permutation analysis. We shuffle local field potential frequencies between recorded units and calculate the amount of phase orders predicted correctly. 25,2% of units exhibited the predicted phase order, significantly more than expected by chance (N=87, permutation test using shuffled labels between frequency / ordering, p<0.05)

